# Coupling of secondary metabolite production in *Bacillus subtilis*

**DOI:** 10.1101/2024.08.15.608099

**Authors:** Caja Dinesen, Manca Vertot, Scott A. Jarmusch, Carlos N. Lozano-Andrade, Aaron J.C. Andersen, Ákos T. Kovács

## Abstract

Although not essential for their growth, the production of secondary metabolites increases the fitness of the producing microorganisms in their natural habitat by enhancing establishment, competition and nutrient acquisition. The Gram-positive soil-dwelling bacterium, *Bacillus subtilis* produces a variety of secondary metabolites. Here, we investigated the regulatory relationship between the non-ribosomal peptide surfactin and the sactipeptide bacteriocin subtilosin A. We discovered that *B. subtilis* mutants lacking surfactin production exhibited higher production of subtilosin A compared to their parental wild-type strain. Additionally, spatial visualization of *B. subtilis* production of metabolites demonstrated that surfactin secreted by a wild-type colony could suppress subtilosin A production in an adjacent mutant colony lacking surfactin production. Reporter assays were performed using mutants in specific transcriptional regulators that confirmed the role of ResD as an activator of the subtilosin A encoding BGC, while removal or Rok and AbrB repressors increased expression of the BGC that was further enhanced by additional deletion of surfactin, suggesting that a so far unidentified regulator might mediate the influence of surfactin on production of subtilosin A. Our study reveals a regulatory influence of one secondary metabolite on another, highlighting that the function of secondary metabolites could be more complex than its influence on other organisms and interactions among secondary metabolites could also contribute to their ecological significance.

**Importance:** Secondary metabolites play an important role in the life of microorganisms facilitating their fitness in the environment, including competing against other microorganisms, interacting with their host or environment, and allowing expansion in their environment. However, secondary metabolites also function as cue molecules influencing gene expression between and within species. Here, we describe that the non-ribosomally synthesized peptide surfactin repress the production of ribosomally synthesized and post translationally modified peptide, subtilosin A in *Bacillus subtilis*, revealing an ecological interaction between two secondary metabolites that could potentially influence the biocontrol efficiency of *B. subtilis* strain that depends on the production of these secondary metabolites against plant pathogen microorganisms.

## Introduction

Biosynthetic gene clusters (BGCs) that are involved in secondary metabolite (SM) production are prevalent across bacterial genera (1, 2). While the production of SMs may not be essential in laboratory settings (3), they likely play a crucial role in the establishment of bacteria within natural niches (4, 5). In the past, the role of SMs in nature has predominately been classified as microbial weapons, likely due to the industrial use of SMs to combat microbial infections (6–8). However, in recent years, this notion has been adjusted. While the antimicrobial properties of SMs are still acknowledged, more research is being directed toward understanding their ecological function rather than being a direct inhibitor of cellular processes (9–12). The soil-dwelling, plant growth-promoting bacterium, *Bacillus subtilis* harbors a diverse array of BGCs, with surfactin and plipastatin being the most studied non-ribosomal lipopeptides (13–15). Particularly, surfactin has a strong biosurfactant activity in addition to its antimicrobial properties (16–18). Surfactin facilitates *B. subtilis* motility through swarming and sliding, thereby playing an important role in *B. subtilis* root colonization in soil (19–21). In addition to non-ribosomally synthesized peptides (NRPs), *B. subtilis* also produces ribosomally synthesized and post-translationally modified peptides (RiPPs). One of the *B. subtilis* specific RiPPs, the bacteriocin subtilosin A was first isolated in 1985 (22) and it displays antibacterial activity towards both Gram-positive and Gram-negative bacteria (22–24). Other functions of subtilosin A have been reported such as suppression of biofilm formation in *Listeria monocytogenes, Gardnerella vaginalis*, and *Escherichia coli* (25). Furthermore, Schoenborn et al. found delayed sporulation in a mutant lacking subtilosin A compared to it parental wild type strain (14).

Whereas surfactin production has been extensively studied across a plethora of *B. subtilis* isolates, research on the production of subtilosin A has predominantly concentrated on the domesticated *B. subtilis* 168 strain or its derivative JH642 (22, 23, 26). Domesticated *B. subtilis* strains lack surfactin production due to mutation in *sfp* gene (27). Importantly, natural isolates of *B. subtilis* encode the intact BGC for subtilosin A production (BGC^Sbo^) (13) and the presence of this BGC is fully conserved among all isolates of *B. subtilis* (28), nevertheless, the production of subtilosin A has not been reported in undomesticated strains.

The BGC^Sbo^, the *sbo-alb* operon encodes the proteins SboA, SboX, and AlbA-AlbG proteins involved in post-translational modifications, processing, and export of the peptide, respectively (23). The BGC^Sbo^ is regulated by several transcription factors, including Rok, AbrB, and ResD, in addition to the sigma factor SigA. Rok and AbrB repress, while ResD activates the expression of the BGC^Sbo^ (29). Production of subtilosin A is linked to later growth stages, characterized by nutrient starvation and oxygen limitation (14, 30). Nakano and colleagues demonstrated that the two-component response regulator, ResDE is essential for activating the subtilosin A BGC in response to oxygen limitation (31).

Several starvation or stationary phase-specific genes are repressed during exponential growth by AbrB, which directly binds to the respective promotors of those genes, as demonstrated for the *sboA* gene. AbrB-mediated repression is alleviated by Spo0A during starvation (32). Additionally, AbrB also repress the transcription of *rok* gene (33, 34). Similarly, Rok binds directly to the promotor of *sboA* and represses its expression (34). While no specific signal or environmental condition has been correlated with the activity of Rok, it is noteworthy that sRok, an interaction partner of Rok, exhibits altered binding affinity during salt stress (35). sRok and DnaA, another interaction partner of Rok, affect the binding affinity of Rok, which may affect Roks regulatory role (36). Moreover, Rok regulates several genes (34, 37), including *sboA,* as well as the biofilm gene *bslA* (*yuaB*) in *B. subtilis* (38). While these studies provide detailed molecular insights into the transcriptional regulation of BGC^Sbo^, the regulation and production of subtilosin A have only been explored in *B. subtilis* 168 and its derivatives that lacks surfactin production, excluding any insights into potential co-dependencies or conflicting expression related to subtilosin A and surfactin production.

In this study, we demonstrate that while the two SMs, surfactin and subtilosin A, are not produced simultaneously, the presence of surfactin regulates the production of subtilosin A in *B. subtilis.* Additionally, we investigate the regulatory mechanism by which surfactin suppresses the expression of the BGC^Sbo^ using knockout mutants in gene encoding transcriptional regulators. Employing GFP reporter assays, analytical chemistry and spatial detection of SMs, we demonstrate that extracellular surfactin inhibits the production of subtilosin A in mutants that otherwise lack surfactin production.

## Results

### Surfactin mutant reveals different production of subtilosin A

Investigation of SM production in the natural isolate *B. subtilis* P8_B1 and its derivative NRP-related BGC mutants using liquid chromatography–mass spectrometry (LC-MS) revealed a varying presence of subtilosin A between P8_B1 and mutants (Fig. 1A). The chemical extractions originating from the mutant derivatives lacking surfactin production (*ΔsrfAC* and *Δsfp*) displayed an additional LC-MS peak corresponding to a m/z of 1134.1963 ([M+3H]^3+^) that was identified as subtilosin A. The same peak is observed in the LC-MS profiles of other isolates that corresponds to mutants lacking surfactin production (13). To confirm this observation in the most frequently used undomesticated *B. subtilis* strain (DK1042 the naturally competent derivative of NCIB3610), samples were extracted from strains DK1042 and Δ*srfAC* to quantify the level of subtilosin A using the peak area. This approach showed an 8.7-fold increase between the peak area in the mutant strain compared to the wild-type DK1042 (P = 0.0193, t-student) (Fig. 1B).

**Fig. 1.**
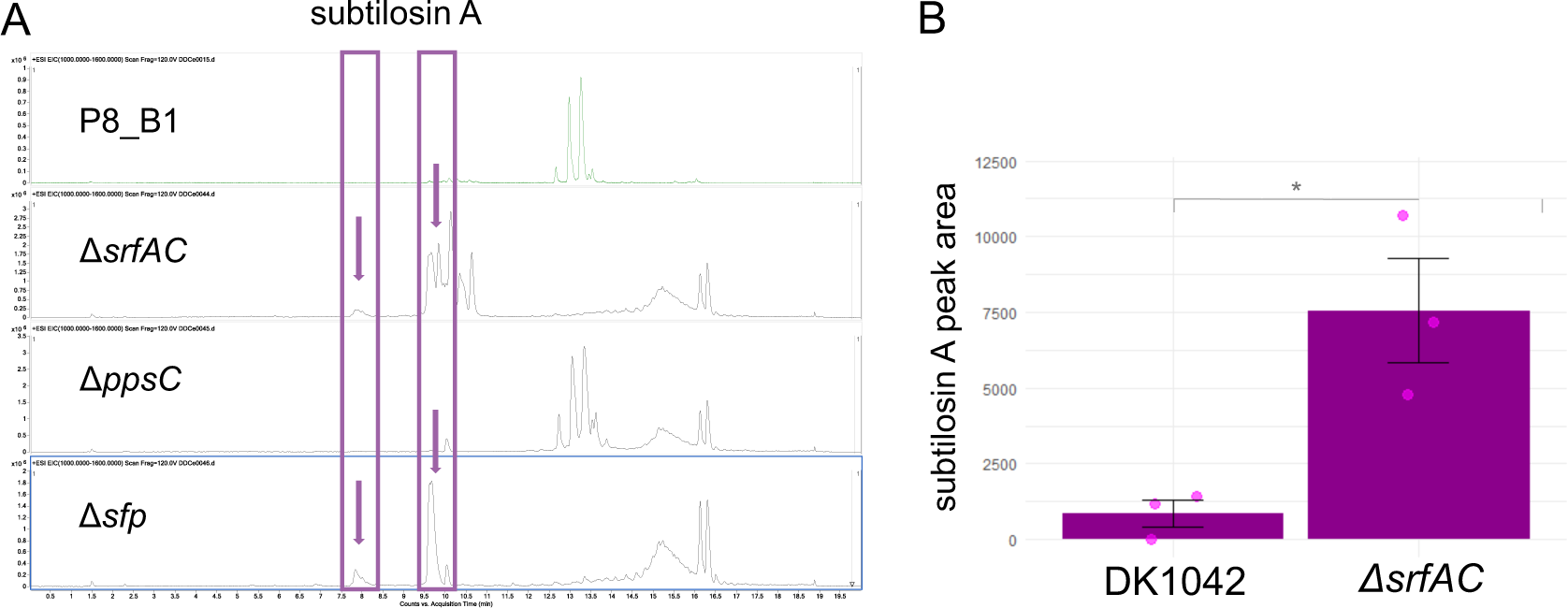
(A) LC-MS chromatogram (EIC: m/z 1000-1600) for *B. subtilis* P8_B1 and its derivative mutants Δ*srfAC* (lacking surfactin), Δ*ppsC* (lacking plipastatin) and Δ*sfp* (lacking all NRPs). Subtilosin Ás peak (1134.1963 [M+3H]3+) is highlighted in the purple boxes. (B) The production of subtilosin A in DK1042 and Δ*srfAC* estimated by peak area from EIC data, statistical difference was tested using students t-test (p = 0.0193, t-student, n=3).

### Surfactin attenuates the expression of BGC^Sbo^

To determine whether the lack of subtilosin A in LC-MS samples from surfactin producers, was due to differentiated production or degradation of subtilosin A, we tested the expression of BGC^Sbo^ in the wild type and mutant derivatives using the GFP signal normalized by OD_600nm_ as proxy. Here, the expression of the BGC^Sbo^ was increased in both the *ΔsrfAC* and the *Δsfp* strains compared to the wild type (p=<0.0001, p=<0.0001, ANOVA and Tukey HSD) (Fig. 2A). To evaluate whether the influence of the lack of surfactin production can be extracellularly complemented, commercially available purified surfactin was supplemented to the Δ*srfAC* strain in varying concentration, showing a reduction in expression of the BGC^Sbo^ with increasing concentration of surfactin (Fig. 2B). Externally added surfactin of 400 µg×ml^-1^ almost reduced the BGC^Sbo^ expression level in the Δ*srfAC* to the levels observed in the wild type (p= 0.8196, ANOVA and Tukey HSD).

**Fig. 2.**
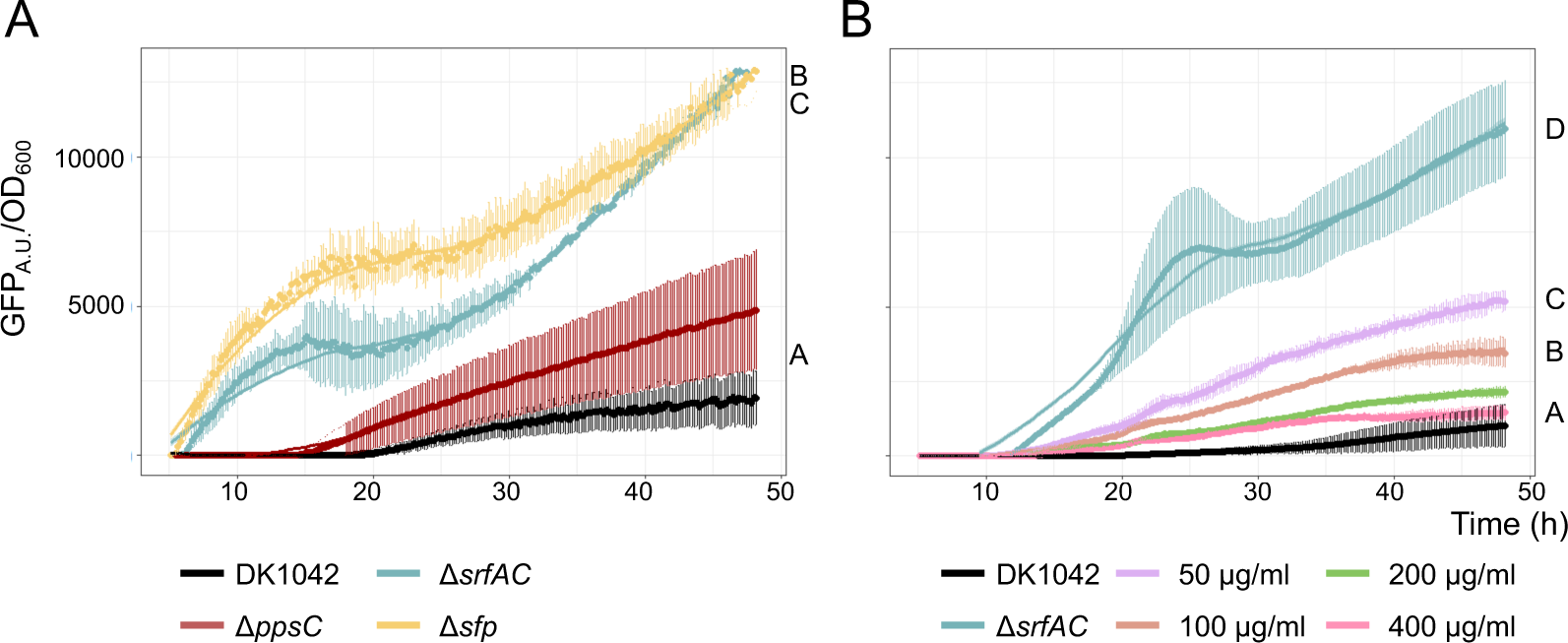
(A) Expression of BGC^Sbo^ in wild type and derived BGC mutants compared using corresponding strains carrying P*_sboA_*-*gfp* reporter fusion. The fluorescence was normalized by growth (optical density at 600nm, OD_600_). (B) Expression of BGC^Sbo^ in Δ*srfAC* strain carrying P*_sboA_*-*gfp* reporter fusion supplemented with varying concentrations of surfactin (50 to 400 µg**×**ml^-1^). Normalized GFP expression between different strains and treatments was compared using the area under the curve (AUC) using one-way ANOVA and Tukey honest test, letters present significant difference between strains (Table S1).

### Complementation of diminished surfactin production in the Δ*srfAC* mutant colony by a neighboring wild-type colony

As external complementation with surfactin can reduce the BGC^Sbo^ expression in the Δ*srfAC* strain similar to the levels seen in the wild type, we investigated whether surfactin production by a wild-type colony could downregulate the expression of BGC^Sbo^ in a neighboring Δ*srfAC* colony. Wild type and Δ*srfAC* strains were spotted next to each other on potato dextrose agar (PDA) medium and sampled for visual detection of SMs. Spatial mapping of metabolites allowed visualization of surfactin production and secretion into the agar by the wild-type strain reaching the proximal edge of the Δ*srfAC* colony. Subtilosin A was detected in a reverse distribution, with high abundance on the distal part of the Δ*srfAC* colony (zone I) with a gradual decrease towards the wild type neighboring edge of the colony (zone II) (Fig. 3AB). Samples were harvested in a line crossing the middle of both colonies (I – IV) and subjected to semi-quantitative LC-MS analysis that verified the diffusion of surfactin from the wild type strain in its environment, in addition to gradually decreasing subtilosin A levels in the Δ*srfAC* colony at increasing surfactin concentrations (Fig. 3C). Additionally, analysis of the spatial metabolite distribution also revealed that production of the sporulation killing factor (SKF) was absent in the Δ*srfAC* colony while abundant in the wild type strain (Fig S1).

**Fig. 3.**
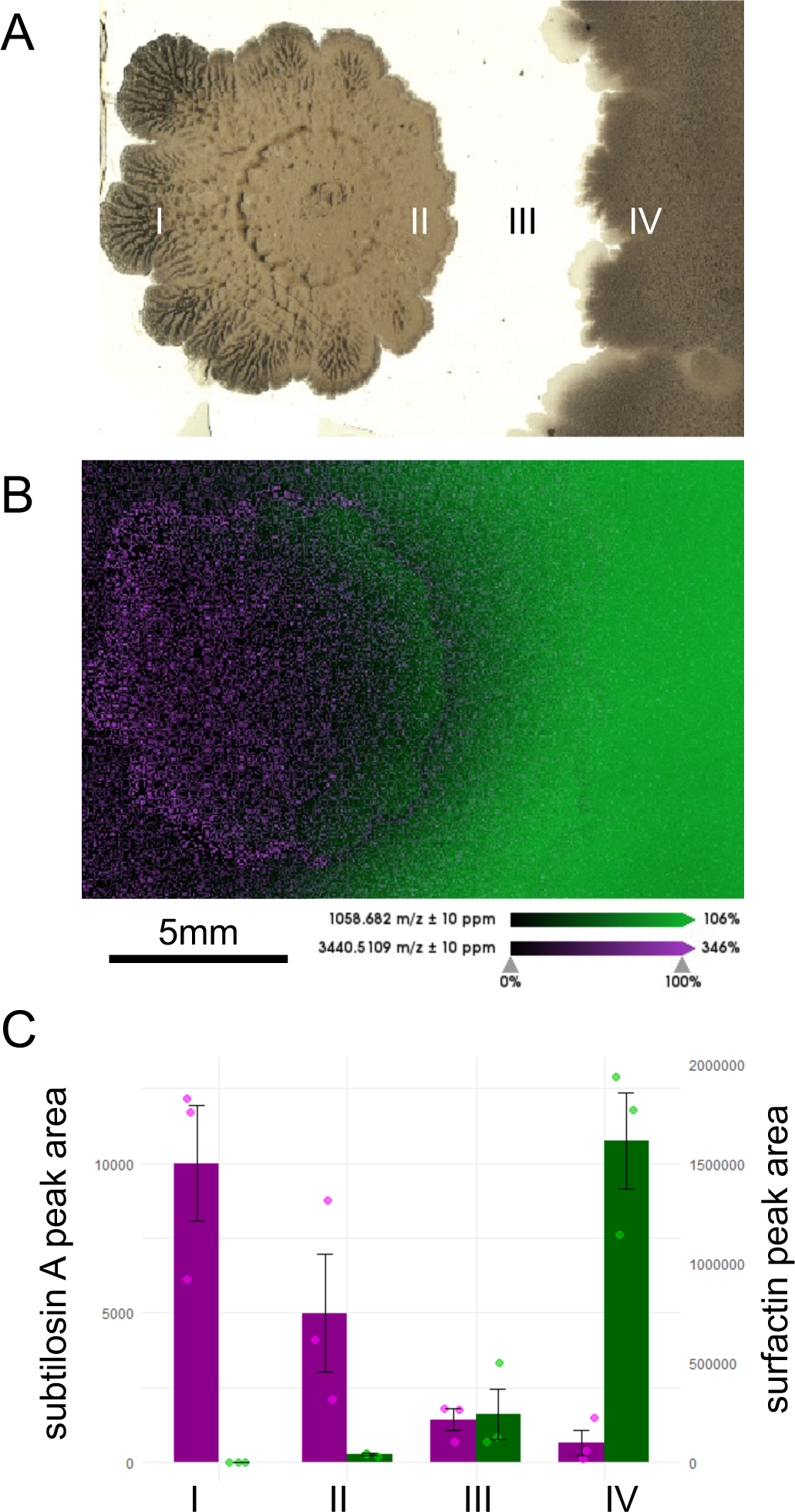
Spatial mapping of subtilosin A and surfactin distribution in neighboring colonies. (A) Light image of Δ*srfAC* (left) and wild type (right) colonies, including the approximate positions of samples taken for LC-MS analysis (I-IV) on a replicate. Scalebar indicates 5 mm. (B) MALDI mass spectrometry imaging based localization of subtilosin A (magenta) and surfactin (green) in the neighboring colonies of Δ*srfAC* and wild-type strains. (C) Relative amount of subtilosin A (magenta) and surfactin (green) estimated by peak area from the LC-MS EIC data in the samples taken at the different positions depicted in panel A.

### Influence of lack of surfactin production on regulation of BGC^Sbo^ by known global regulators

To evaluate whether surfactin downregulates BGC^Sbo^ expression through one of the known global regulators of BGC^Sbo^, *resD*, *rok*, and *abrB* genes were disrupted in the wild type and *ΔsrfAC* carrying the P*_sboA_*-*gfp*. Deletion of *resD* prevented the expression of *sboA* (Fig. 4A), whereas introduction of Δ*rok* and Δ*abrB* increased expression of BGC^Sbo^ (p = <0.0001 and p = <0.0001, ANOVA and Tukey HSD) (Fig. 4BC). The combination of Δ*srfAC* with Δ*resD* did not influence the already diminished expression of BGC^Sbo^ (Fig. 4A). In contrast, deletion of the BGC for surfactin production in the Δ*rok* further increased the expression level of BGC^Sbo^ (p = <0.0001, ANOVA and Tukey HSD) (Fig. 4BC). While in the absence of *rok* gene, this increase was maintained throughout the experiment, BGC^Sbo^ expression in the Δ*abrB* Δ*srfAC* mutant was only enhanced in the first 20 h, whereas afterward was comparable to the single Δ*abrB* strain with no statistical difference (p = 0.5463, ANOVA and Tukey HSD) (Fig. 4C).

**Fig. 4.**
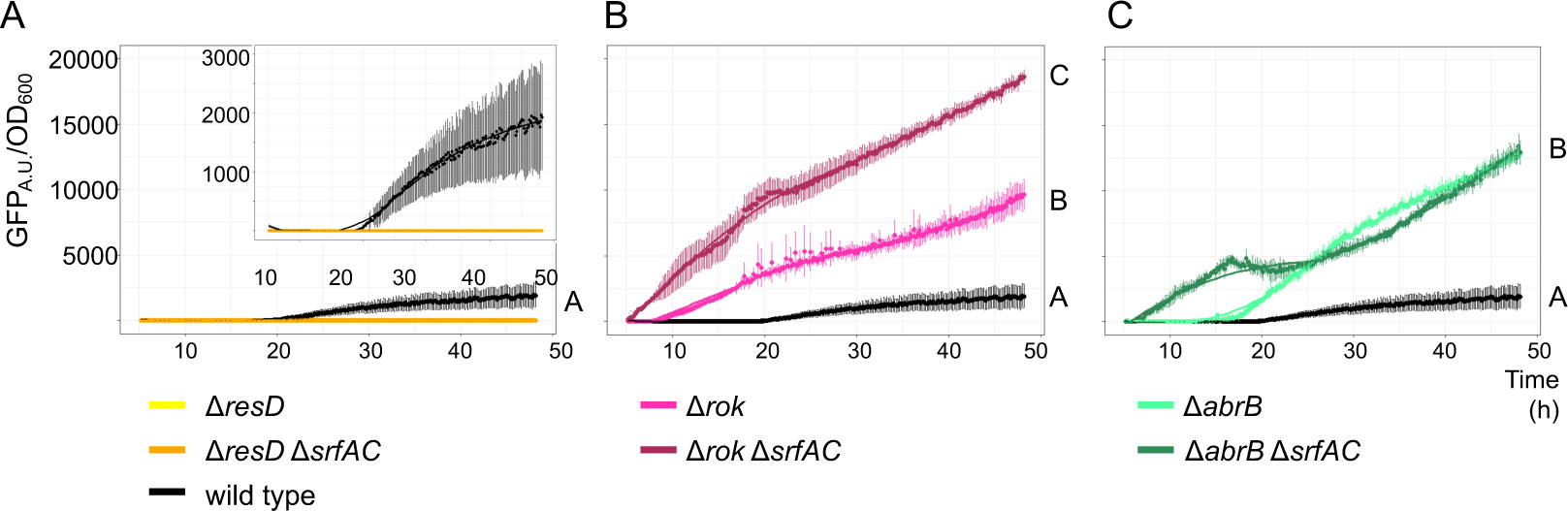
Expression of BGC^Sbo^ in wild type (black line) and derived regulator mutants compared using corresponding strains carrying P*_sboA_*-*gfp* reporter fusion. The fluorescence was normalized by growth (optical density at 600 nm, OD_600_). Expression was assayed in Δ*resD* (A), Δ*rok* (B), and Δ*abrB* (C) single mutants (light line colors) or in combination with Δ*srfAC* (dark line colors). Normalized GFP expression between different strains and treatments was compared using the area under the curve (AUC) using one-way ANOVA and Tukey honest test, letters represent significant difference between strains (Table S2).

## Discussion

SMs have been extensively investigated and harnessed, playing a pivotal role in combating microbial infections and improving human health (4, 5), with a growing interest in the application of SMs beyond medicine (6, 7). In particular, understanding the ecological functions of SMs can enhance the utilization of SM producing bacteria in agricultural applications, where production of SM is important for the efficiency of biocontrol bacteria. Identification of the underlying regulatory mechanisms influencing SM production may facilitate elucidating their role in nature.

Here, we dissected the influence of the lipopeptide surfactin on expression and production of the bacteriocin subtilosin A in *B. subtilis*. Surfactin decreased the production of subtilosin A in *B. subtilis*, while strains lacking surfactin production had increased level of subtilosin A. The lack of surfactin production and therefore enhanced subtilosin A level could be reverted by pure surfactin or inoculating a neighboring wild-type colony next to the Δ*srfAC* mutant strain. Testing the expression of BGC^Sbo^ demonstrated a transcriptionally regulatory mechanism behind surfactin-mediated repression of subtilosin A production. Previous studies have demonstrated both overlapping and dissimilar production of SMs in *B. subtilis* (39). For example, Yannarell et al. reported little overlap of cells expressing both BGCs for surfactin and subtilosin A production in biofilm colonies (39). Spatial detection of key SMs in *B. subtilis* biofilm colonies has been previously reported using MALDI-MSI (40). Although not specifically reported, the MALDI-MSI images suggest increased subtilosin A production in the Δ*srfAA* mutant colony confirming our results. Similarly, reduced SKF level was noticeable in the Δ*srfAA* mutant used by Si and colleagues (40), which again confirms our data.

The lack of simultaneous production of surfactin and subtilosin A might suggest that their roles in *B. subtilis* are distinctive and these SMs might contribute to different developmental stages or specific environmental conditions. Or rather, that in the absence of surfactin, subtilosin A antibacterial properties are replacing that of surfactins. While experimental validation is required to demonstrate such possibility, various roles of RIPPs have previously been reported, such as growth inhibition, nutrient competition and quorum sensing (41). Notably, surfactin plays a pivotal role in the early stages of root colonization in soil and during initiation of biofilm formation (19–21), congruent with the early exponential phase expression of the *srfA* operon, around 7 hours after inoculation (42). On the contrary, production of subtilosin A is correlated with the end of the exponential phase/starting stationary phase (30). The quantities in which *B. subtilis* produce these SMs is also different. The level of surfactin has been quantified in different *B. subtilis* strains, ranging from 1.25 – 6.45 g×l^-1^ (43–46), while subtilosin A concentration in different strains and conditions has been reported to be between 0.5 – 7.8 mg×l^-1^ (22, 30). The production of surfactin and subtilosin A was not measured quantitatively in our study; however, the LC-MS data suggest that surfactin was produced in higher quantities than subtilosin A. The difference in production quantity might be related to their role in the environment, since the function of surfactin as a bio-surfactant may require higher quantities compared to the primarily antibiotic role of subtilosin A.

The gene cluster encoding subtilosin A synthesis is known to be regulated by the global regulators ResD, Rok, and ArbB (29). Our analysis with *sboA* promoter coupled *gfp* reporter strains confirmed current knowledge on the role of ResD, Rok, and AbrB in the transcriptional regulation of BGC^Sbo^ (29). Disruption of surfactin production further increased the expression of BGC^Sbo^ on a Δ*rok* background, suggesting that Rok is not involved in perceiving the presence of surfactin. Since ResD works as an activator of BGC^Sbo^ expression, deletion of both *resD* and *srfAC* does not permit the demonstration whether surfactin influences ResD. AbrB functions as a repressor of BGC^Sbo^ transcription, with its repression being relieved during starvation. While disruption of surfactin production in a Δ*abrB* background hastened the expression of BGC^Sbo^in the first 20h compared to the single deletion of *abrB*, the expression levels of BGC^Sbo^ were comparable in the two strains from 20h onwards. The enhanced expression of BGC^Sbo^ observed in the earlier phase of the population growth in a Δ*abrB* background, when expression of surfactin related BGC is prominent, suggests that surfactin is not regulating subtilosin A production through AbrB. Interestingly, while deletion of either *srfAC*, *sfp*, or *rok* increases BGC^Sbo^ expression from the first few hours of the population growth, disruption of *abrB* only influences BGC^Sbo^ expression 20 h after inoculation of the culture. These experiments suggest an additional regulatory system might be involved in perceiving the presence of surfactin in *B. subtilis*. Examination of the wild-type and Δ*srfAC B. subtilis* transcriptome could potentially reveal which genes and regulatory pathways are primarily influenced by surfactin. This could additionally reveal if the transcription of BGCs other than BGC^Sbo^, are differentially regulated in the absence of surfactin, in accordance with the decreased SKF level detected in the Δ*srfAC* mutant colony.

Identifying possible correlations and differences in the production of SMs in *B. subtilis*, such as that described here between subtilosin A and surfactin, could further increase our understanding of the ecological roles of SMs.

## Materials and methods

### Bacterial strains and culture media

All strains used in this study, including genomic DNA (gDNA) donors, are listed in Table 1. Overnight starter cultures were grown in lysogeny broth (LB, Carl Roth, Germany; 10 g×l^−1^ tryptone, 5 g×l^−1^ yeast extract, and 5 g×l^−1^ NaCl) medium. If not stated otherwise, experiments were performed in potato dextrose broth (PDB; BD, USA; potato infusion at 4 g×l^−1^, glucose at 20 g×l^−1^), supplemented with 1.5 % agar when required.

**Table 1.** Detailed information about strains used in this study.

| Strain | Description | Reference |
| --- | --- | --- |
| 168 | *amyE::P<sub>sboA</sub>-gfp* (Cm<sup>R</sup>) | (47) |
| DK1042 | NCIB 3610 *comI*<sup>Q12</sup> | (48) |
| DS1122 | 3610 Δ*srfAC* (MIs<sup>R</sup>) | (49) |
| DS4114 | 3610 Δ*ppsC* (Tet<sup>R</sup>) | (50) |
| DS3337 | 3610 Δ*sfp* (MIs<sup>R</sup>) | (51) |
| P8_B1 | WT | (13) |
| P8_B1 | Δ*srfAC* (MIs<sup>R</sup>) | (13) |
| P8_B1 | Δ*ppsC* (Tet<sup>R</sup>) | (13) |
| P8_B1 | Δ*sfp* (MIs<sup>R</sup>) | (13) |
| DTUB366 | DK1042 *amyE::P<sub>sboA</sub>-gfp* (Chl<sup>R</sup>) | This study |
| DTUB367 | DK1042 *amyE::P<sub>sboA</sub>-gfp* (Chl<sup>R</sup>); Δ*srfAC* (MIs<sup>R</sup>) | |
| DTUB368 | DK1042 *amyE::P<sub>sboA</sub>-gfp* (Chl<sup>R</sup>); Δ*ppsC* (Tet<sup>R</sup>) | |
| DTUB369 | DK1042 *amyE::P<sub>sboA</sub>-gfp* (Chl<sup>R</sup>); Δ*sfp* (MIs<sup>R</sup>) | |
| DTUB370 | DK1042 *amyE::P<sub>sboA</sub>-gfp* (Chl<sup>R</sup>); Δ*rok* (Km<sup>R</sup>) | |
| DTUB371 | DK1042 *amyE::P<sub>sboA</sub>-gfp* (Chl<sup>R</sup>); Δ*resD* (Km<sup>R</sup>) | |
| DTUB372 | DK1042 *amyE::P<sub>sboA</sub>-gfp* (Chl<sup>R</sup>); Δ*abrB* (Km<sup>R</sup>) | |
| DTUB373 | DK1042 *amyE::P<sub>sboA</sub>-gfp* (Chl<sup>R</sup>); Δ*rok* (Km<sup>R</sup>), Δ*srfAC* (MIs<sup>R</sup>) | |
| DTUB374 | DK1042 *amyE::P<sub>sboA</sub>-gfp* (Chl<sup>R</sup>); Δ*resD* (Km<sup>R</sup>), Δ*srfAC* (MIs<sup>R</sup>) | |
| DTUB375 | DK1042 *amyE::P<sub>sboA</sub>-gfp* (Chl<sup>R</sup>); Δ*abrB* (Km<sup>R</sup>), Δ*srfAC* (MIs<sup>R</sup>) | |

### Generation of mutant *B. subtilis* strains

DK1042 P*_sboA_-gfp* was obtained with gDNA from the gDNA donor 168 *amyE::*P*_sboA_-gfp*. Mutant strains in DK1042 P*_sboA_-gfp* were obtained by natural competence (52), by transforming gDNA and selecting for antibiotic (AB) resistance on AB containing LB agar medium. gDNA was extracted from the donors using the EURx Bacterial & Yeast Genomic DNA Purification Kit (EURx, Gdansk, Poland), following the manufactureŕs instructions. To verify transformation and lack of SM production, overnight grown cultures were directly extracted with acetonitrile using a 1:1 acetonitrile:culture dilution, where after the solution was centrifuged and supernatant transferred to HPLC vials and analyzed by ultrahigh performance liquid chromatography coupled to high-resolution mass spectrometry (UHPLC-HRMS).

### Expression assay in *B. subtilis* BGC mutants

The effect of SM production on the expression BGC^Sbo^ was evaluated in plate reader assays. Fluorescence and optical density were detected in cultures grown in 96-well microtiter plates with 200 µl PDB including the reporter strains with a final optical density of 0.01 at 600nm (OD_600_). To test the influence of surfactin on the expression of *sboA*, a similar setup was used, except the P*_sboA_*-*gfp* Δ*srfAC* strain was supplemented with surfactin at a final concentration of 50, 100, 200 and 400 µg×ml^-1^. PDB medium without surfactin served as a control. Cultivation was performed in Synergy XHT multi-mode reader (Biotek Instruments, Winooski, VT, US), at 30°C with orbital continuous shaking (3 mm), monitoring the OD_600_ as well as GFP (Ex: 482/20; Em:528/20; Gain: 60) fluorescence every 5 min.

### Detection of subtilosin A and surfactin in neighboring colonies of wild-type and Δ*srfAC* strains

Complementation of surfactin production by the wild type colony towards the neighboring ΔsrfAC mutant was tested on PDA medium. 2 µl overnight grown bacterial cultures were inoculated on PDA medium using a 2.5 cm distance between the inoculation points of the two strains. The plates were incubated at 37oC for 3 days. To assess the level of surfactin and subtilosin A, four plugs were transferred from the plates distributed from the distal region of the ΔsrfAC colony to the distal edge of the wild-type colony (see Fig. 3). 1.5 ml isopropanyl:ethyl acetate (1.3 v/v) with 1% formic acid was added to each plug and sonicated for 60 min before centrifugation (3 min, 13400 rpm). The supernatant was extracted and transferred under N2 with no heat before resuspension in 250 µl methanol and centrifugation (3 min, 13400 rpm). Supernatant was transferred to HPLC vials and tested by UHPLC-HRMS.

UHPLC-HRMS was performed on an Agilent Infinity 1290 UHPLC system with a diode array detector. UV–visible spectra were recorded from 190 to 640 nm. Liquid chromatography of 1 μL extract (or standard solution) was performed using an Agilent Poroshell 120 phenyl-hexyl column (2.1 × 150 mm, 1.9 μm) at 40 °C using acetonitrile (ACN) and H2O, both containing 20 mM formic acid, as mobile phases. Initially, a gradient of 10% ACN/H2O to 100% acetonitrile over 10 min was employed, followed by isocratic elution of 100% ACN for 2 min. The gradient was returned to 10% ACN/H2O in 0.1 min, and finally isocratic condition of 10% ACN/H2O for 2.9 min, at a flow rate of 0.35 ml×min-1. HRMS spectra were acquired in positive ionization mode on an Agilent 6545 QTOF MS equipped with an Agilent Dual Jet Stream electrospray ion source with a drying gas temperature of 250 °C, drying gas flow of 8 l×min-1, sheath gas temperature of 300 °C, and sheath gas flow of 12 l×min^-1^. Capillary voltage was set to 4000 V and nozzle voltage to 500 V. MS data analysis and processing were performed using Agilent MassHunter Qualitative Analysis B.07.00.

### Mass spectrometry imaging of pairwise interactions between Δ*srfAC* and wild-type colonies

Samples were prepared as described above for quantification of SMs from PDA grown colonies. Interaction zone of the two colonies were excised from agar plates and adhered to MALDI IntelliSlides (Bruker, Billerica, Massachusetts, USA) using a 2-Way Glue Pen (Kuretake Co., Ltd, Nara-Shi, Japan). Slides were covered by spraying 1.5 ml of 2,5-dihydrobenzoic acid (40 mg×ml^-1^ in MeOH/H_2_O (80:20, v/v, 0.1% trifluoroacetic acid)) and dried prior to MSI acquisition. MALDI-MSI data was acquired using a timsTOF flex (Bruker Daltonik GmbH) mass spectrometer operating in a positive mode with 30 µm raster width and a m/z range of 500– 4000. Calibration was performed by using red phosphorus. The settings in the timsControl were as follow: Laser: imaging 30 µm, Power Boost 3.0%, scan range 26 µm in the XY interval, and laser power 70%; Tune: Funnel 1 RF 300 Vpp, Funnel 2 RF 300 Vpp, Multipole RF 300 Vpp, isCID 0 eV, Deflection Delta 70 V, MALDI plate offset 100 V, quadrupole ion energy 5 eV, quadrupole loss mass 100 m/z, collision energy 10 eV, focus pre TOF transfer time 75 µs, pre-pulse storage 8 µs. Data was root mean square normalized and visualized in SCiLS software (Bruker Daltonik GmbH, Bremen, Germany).

### Statistics

Data was analyzed and graphically represented using R 4.3.2 and the package ggplot2 (53). Student’s t-test was used to test for statistical differences in experiments with two groups. Statistical significance (α) was set at 0.05. For multiple comparisons (more than two treatments), one-way analysis of variance (ANOVA) and Tukey’s honestly significant difference (HSD) were performed. In all the cases, normality and equal variance were assessed using the Shapiro - Wilks and Levene test, respectively.

## Acknowledgement

This project was supported by Novo Nordisk Foundation via the project INTERACT (NNF19SA0059360). Funding from the Danish National Research Foundation (DNRF137) for the Center for Microbial Secondary Metabolites and Novo Nordisk Foundation for the infrastructure “Imaging microbial language in biocontrol (IMLiB)” (NNFOC0055625) is acknowledged. The Metabolomics Core (DTUMetCore) of the Technical University of Denmark’s Bioengineering Department is acknowledged for access to analytical instrumentation.

## Author contributions

Designed research: CD, ÁTK; performed the experiments: CD; analysis of strains: CNLA; performed the chemical detection and analysis: MV, SJ, AJCA; analyzed data: CD; wrote the manuscript: CNLA, ÁTK with corrections by co-authors.

## Supplementary information

**Fig. S1.**
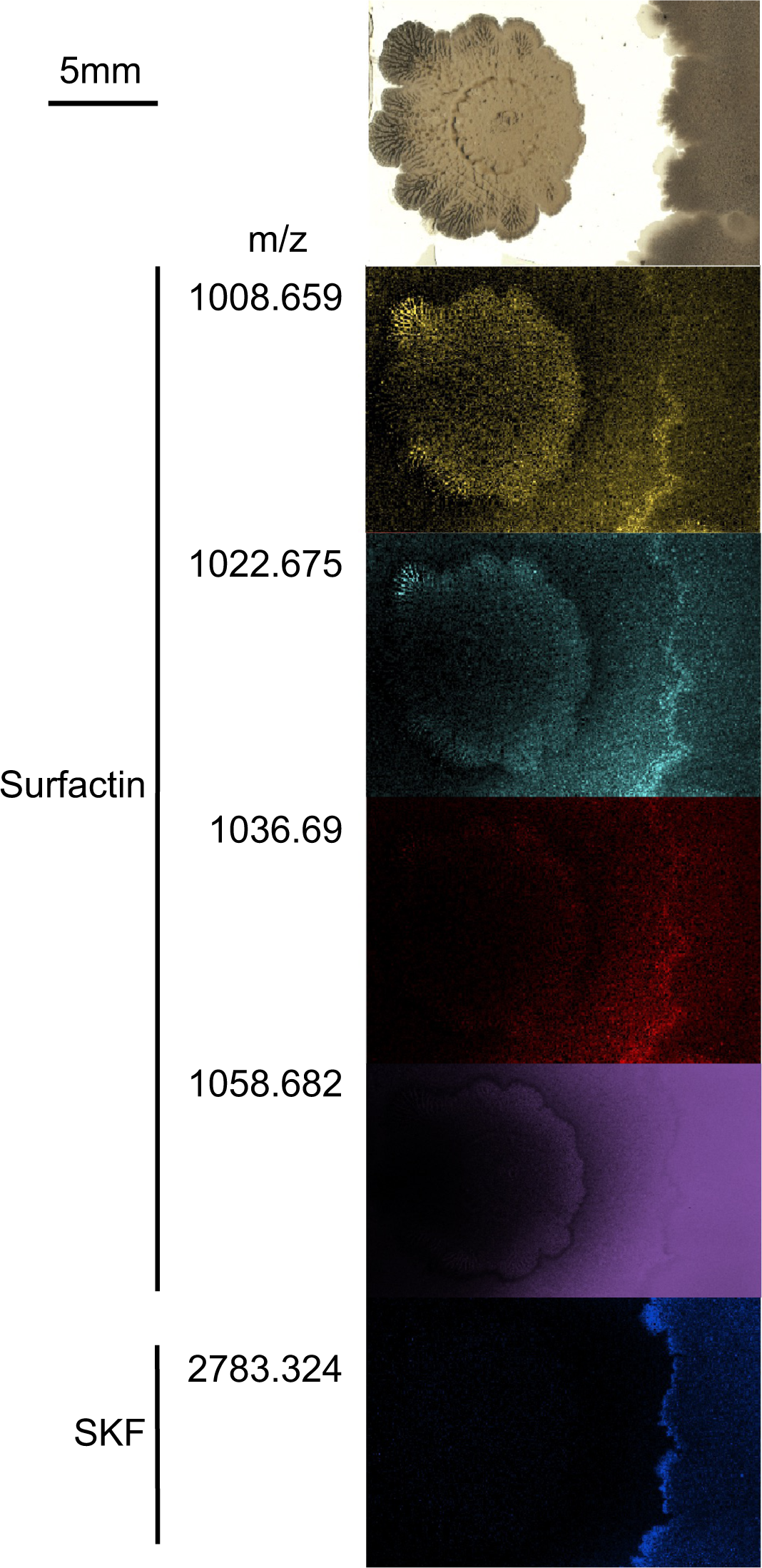
Spatial mapping of surfactin isomer and SKF distribution in neighboring colonies. Top row includes the light image of Δ*srfAC* (left) and wild type (right) colonies. Scalebar indicates 5 mm. MALDI mass spectrometry imaging-based localization of surfactin isomers and SKF in the neighboring colonies of Δ*srfAC* and wild-type strains.

**Table S1.** Statistics on data used in Fig 2.

| Fig 2A |  | Fig 2B |  |
| --- | --- | --- | --- |
| WT – *srfAC* **** | p < 0.0001 | WT – *srfAC* **** | p < 0.0001 |
| WT – *ppsC* ns | p = 0.115 | WT – 50 *** | p = 0.0003 |
| WT – *sfp* **** | p < 0.0001 | WT – 100 * | p = 0.0182 |
| *srfAC* – *sfp* * | p = 0.0492 | WT – 200 ns | p = 0.4597 |
|  |  | WT – 400 ns | p = 0.8196 |

**Table S2.** Statistics on data used in Fig 4.

| Fig 4A |  |
| --- | --- |
| WT – *resD* | ns |
| WT – *resD*, *srfAC* | ns |
| *resD* – *resD*, *srfAC* | ns |
| Fig 4B |  |
| WT – *rok* **** | p < 0.0001 |
| WT – *rok*, *srfAC* **** | p < 0.0001 |
| *rok* – *rok*, *srfAC* **** | p < 0.0001 |
| Fig 4C |  |
| WT – *abrB* **** | p < 0.0001 |
| WT – *abrB*, *srfAC* **** | p < 0.0001 |
| *abrB* – *abrB*, *srfAC* ns | p = 0.5463 |

